# *Drosophila* carboxypeptidase D (SILVER) is a key enzyme in neuropeptide processing required to maintain locomotor activity levels and survival rate

**DOI:** 10.1101/551853

**Authors:** Dennis Pauls, Yasin Hamarat, Luisa Trufasu, Tim M. Schendzielorz, Gertrud Gramlich, Jörg Kahnt, Jens T. Vanselow, Andreas Schlosser, Christian Wegener

## Abstract

Neuropeptides are processed from larger preproproteins by a dedicated set of enzymes. The molecular and biochemical mechanisms underlying preproprotein processing and the functional importance of processing enzymes are well characterised in mammals, but little studied outside this group. In contrast to mammals, *Drosophila* lacks a gene for carboxypeptidase E (CPE), a key enzyme for mammalian peptide processing.

By combining peptidomics and neurogenetics, we addressed the role of *Drosophila* carboxypeptidase D (dCPD) in global neuropeptide processing and selected peptide-regulated behaviours. We found that a deficiency in dCPD results in C-terminally extended peptides across the peptidome, suggesting that dCPD took over CPE function in the fruit fly. dCPD is widely expressed throughout the nervous system, including peptidergic neurons in the mushroom body and neuroendocrine cells expressing adipokinetic hormone. Conditional hypomorphic mutation in the dCPD-encoding gene *silver* in the larva causes lethality, and leads to deficits in adult starvation-induced hyperactivity and appetitive gustatory preference, as well as to reduced survival rate and activity levels. A phylogenomic analysis suggests that loss of CPE is not a common insect feature, but specifically occured in Hymenoptera and Diptera. Our results show that dCPD is a key enzyme for neuropeptide processing in *Drosophila*, and is required for proper peptide-regulated behaviour. dCPD thus appears as a suitable target to genetically shut down total neuropeptide production in peptidergic neurons. Our results raise the question why *Drosophila* and other Diptera and Hymenoptera –unlike other insects-have obviously lost the gene for CPE but kept a gene encoding CPD.

## Introduction

Neuropeptides and peptide hormones are synthesised as parts of larger precursors (preproproteins) from which they are processed into their bioactive form by sequential action of proprotein convertases (PCs), metallocarboxypeptidases (CPs) and amidating enzymes (1–3). In mammals, proprotein processing is well characterised on the molecular and biochemical level, and is implicated in a variety of physiological and pathological processes including obesity and growth defects (4, 5). In contrast, the mechanisms and functions of proprotein processing are little studied in lower vertebrate taxa and invertebrates.

For insects, preproprotein processing is best understood for *Drosophila* (reviewed by (6)). In the fruit fly, the first processing step is catalysed by the proprotein convertase dPC2 encoded by the gene *amontillado* (*amon*) (7–9). A deficiency in dPC2 results in reduced to absent levels of neuro- and enteroendocrine peptides (9–11), developmental defects and impaired behaviour including hatching and ecdysis (7, 8, 12) and larval locomotion (9) as well as carbohydrate metabolism (10). As a typical PC, dPC2 cuts C-terminal of canonic mono- or dibasic amino acid (typically R, KR or RR) cleavage sites, resulting in peptides which are C-terminally extended by basic amino acids. In mammals, these basic C-terminal extensions are pruned by N/E metallocarboxypeptidase E (CPE, EC 3.4.17.10 (13–15)), which initially was thought to be the only carboxypeptidase involved in neuropeptide processing (16). However, a further member of the M14 subfamily of metallocarboxypeptidases, carboxypeptidase D (CPD, EC 3.4.17.22), was later found to be able to partially compensate for a loss of CPE action (17). CPDs are unusual in that they are large proteins (∼180 kDa) that contain three CP domains. Only the first two domains are enzymatically active, while the third domain is inactive (16, 18). In contrast, CPEs and other M14 CPs like the membrane-bound CPM (EC 3.4.17.12) are smaller (52-56 kDa) and contain only one CP domain (19, 20).

Curiously, unlike vertebrates and most other invertebrates, *Drosophila* lacks a CPE gene, and only possesses a CPD-encoding gene named *silver* (21). The two active domains of dCPD differ in their pH optima and substrate preferences (22–24). While null mutations in *svr* are lethal (25), dCPD transgenes containing either active domain 1 or 2 can rescue behavioural and developmental deficits of *svr* mutant flies to varying degrees (24). Hypomorphic mutants have defects in cuticle melanisation and tanning, wing morphogenesis, biogenic amine metabolism, light response (26, 27) and memory formation (24, 28), and show increased ethanol and cold sensitivity (24).

The general requirement of CPs in neuropeptide and peptide hormone processing and the lack of a CPE gene in the *Drosophila* genome implies that dCPD is functionally important for the production of bioactive peptides in the fruit fly. This reasoning is supported by a prominent expression of *svr* in the central nervous system (CNS) and endocrine tissues such as the midgut (29), as well as by the direct transactivation of *svr* by the basic helix-loop-helix transcription factor DIMMED which confers a neuroendocrine peptidergic phenotype (30). Ectopic expression of *svr* transgenes in the neurohemal organ of the brain (corpora cardiaca) also affects processing of adipokinetic hormone (AKH) (24). Yet, direct and comprehensive biochemical evidence for a role of dCPD or any other insect CP in neuropeptide processing is lacking.

Here, we used a combined neurogenetic-peptidomic approach to test the requirement of dCPD for neuropeptide processing, locomotor behaviour and life span. A phylogenomic search for CPD and CPE genes throughout the insects suggests that CPE has been independently lost in a few holometabolous insect taxa, while all analysed insect species obviously possess CPD. Taken together, our results suggest that insect CPDs can fully compensate for a loss of CPE and possibly have important further functions beyond neuropeptide processing that cannot be fulfilled by CPE.

## Results

### Characterisation of the transgenic heat-shock rescue lines

The *svr*^*PG33*^ allele is lethal in hemizygous males and homozygous females (25). To test the functionality of the newly generated *hs-svr* rescue lines, we crossed *svr*^*PG33*^/FM7h virgins with males of two newly generated FM7/Y; *hs-svr* lines and kept the flies either at constant 18°C or under a heat-shock regime. Once all offspring had eclosed, we scored the F1 males for the dominant FM7 marker *Bar (B)*. At constant 18°C, only *B* males (FM7/Y; *hs-svr*/+) were present in both crosses (59/40 males, respectively), confirming the lethality of the *svr*^*PG33*^ phenotype. Under heat-shock regime, both *B* and normal-eyed males were found (36:33/37:13, respectively), indicating a ≥35% heat-shock rescue efficiency for *svr*^*PG33*^/Y males. Thus, the newly generated *hs-svr* lines are functional and can be used to rescue lethal mutations in *svr*.

While a *hs-*promoter is ideal to drive rescue gene expression in non-feeding developmental stages (egg, pupa), it is prone to low background expression at normal temperatures (31). Using RT-PCR with *svr-*specific primers (Fig. 1A), we found moderate *svr* expression in *w*^*1118*^ control flies containing a functional endogenous *svr* gene, high *svr*-expression in heatshock-rescued *svr*^*PG33*^, *hs-svr* mutant flies, and low expression in *svr*^*PG33*^, *hs-svr* flies maintained at 18°C (Fig. 1B). To distinguish whether the *svr*-expression in *svr*^*PG33*^, *hs-svr* flies at low temperature is due to background expression from the transgene or expression from the mutated endogenous *svr*^*PG33*^ allele, we performed RT-PCR with a *GawB* insert-specific primer set (Fig. 1C). *GawB* expression was absent from w^1118^ controls, but detectable in heat-shock-rescued *svr*^*PG33*^ mutant flies. Again, low expression was found for *svr*^*PG33*^, *hs-svr* flies maintained at 18°C. A primer set spanning the region from the 3’ end of *GawB* to *svr* exon 5 did not result in a detectable product in all flies and conditions tested (data not shown). This confirms previous findings by northern blots showing that the *PG*^*33*^ insertion disrupts expression of native *svr* mRNA ((23). We thus concluded that *svr*^*PG33*^ mutant flies do not express a functional native dCPD. Instead, the *hs-svr* rescue construct seems to be expressed at low levels even at lower temperatures. Without heat-shock, *svr*^*PG33*^, *hs-svr* flies thus represent functional dCPD hypomorphs.

**Figure 1:**
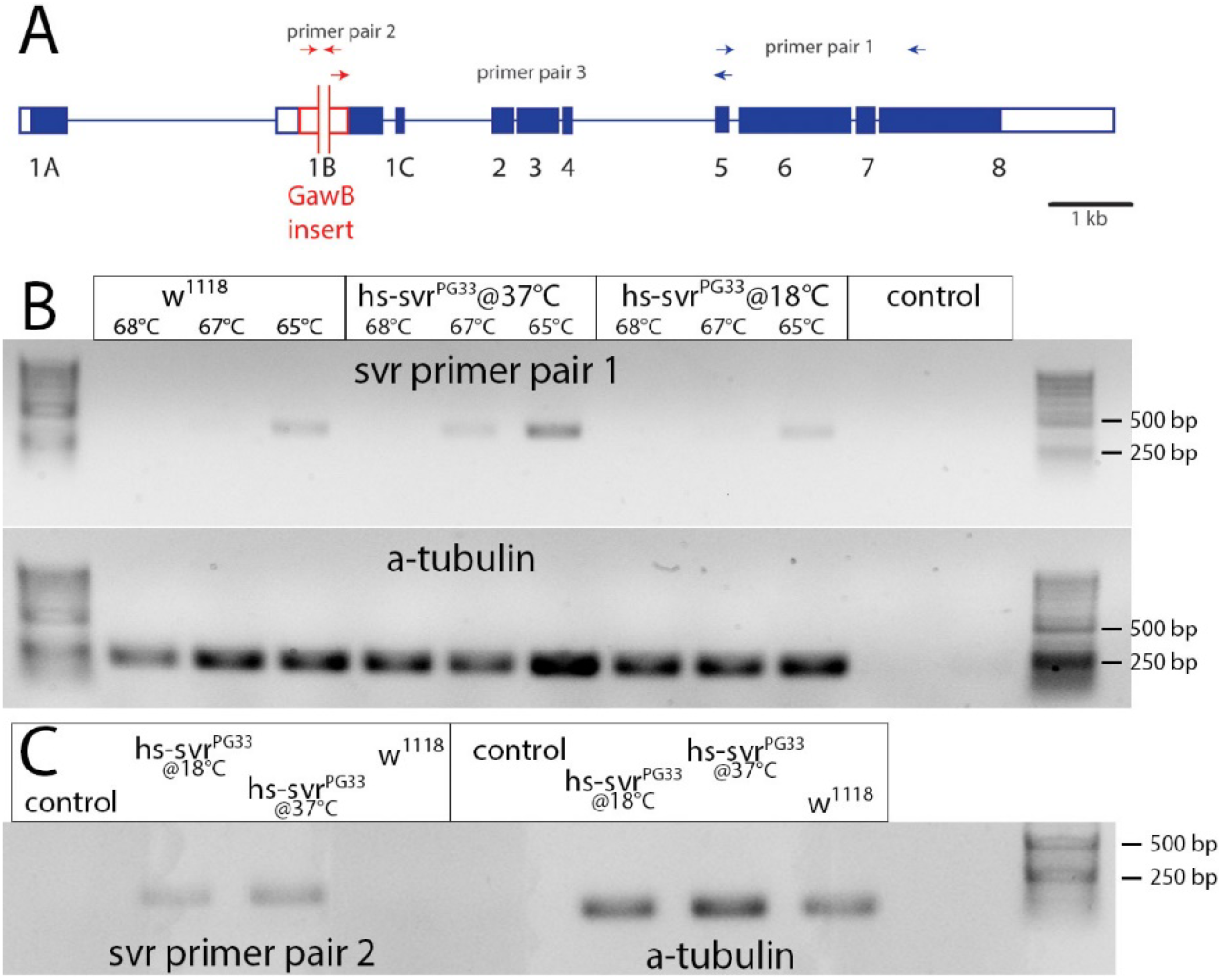
RT-PCR analysis of svr expression in w^1118^ (control) and svr^PG33^;hs-svr (hs-svr^PG33^) flies. **A)** Genomic structure of the svr gene with the svr^PG33^ GawB insert. Primer pairs used for RT-PCR are indicated. Blue boxes represent exons. **B)** Agarose gel electrophoresis of amplificates obtained with primer pair 1 that spans exon 5 to exon 8 at three different annealing temperatures. A band is visible at the optimum annealing temperature (65°C) in w^1118^ flies, indicating that svr is expressed (top left). The same band is much stronger in heat-shocked svr^PG33^;hs-svr flies, indicating heat-shocked induced svr expression from the hs-svr genomic insert (top middle). Without heat-shock, a similar yet much weaker band is visible in svr^PG33^;hs-svr flies (top right), indicating background expression of the svr rescue construct. α-Tubulin was used as standard (bottom). **C)** Agarose gel electrophoresis of amplificates obtained with primer pair 2 that spans part of the GawB insert which disrupts expression of the native svr gene. A band is visible in svr^PG33^;hs-svr flies independent of heat-shock, but is absent in w^1118^ controls (left). α-Tubulin was used as standard (right). Control: sample without cDNA.

### Deficiency of dCPD SILVER results in incompletely processed neuropeptides and peptide hormones

To qualitatively test whether a defective dCPD results in changes in peptide processing, we raised *svr*^*PG33*^/FM7h; *hs-svr* flies under heat-shock conditions until eclosion, followed by fourteen days at constant 18°C. Then, the nervous system of individual males was dissected and either optic lobes (n=15 controls/17 mutants), central brain without optic lobes (n=12/16) or the ventral ganglion (n=14/9) were separated and directly profiled by MALDI-TOF MS (Fig. 2A-C). This method comprises specific on-plate extraction of peptides, which allows to identify peptides by mass match with high reliability in *Drosophila* (32, 33) and other insects (e.g. (34)).

**Figure 2:**
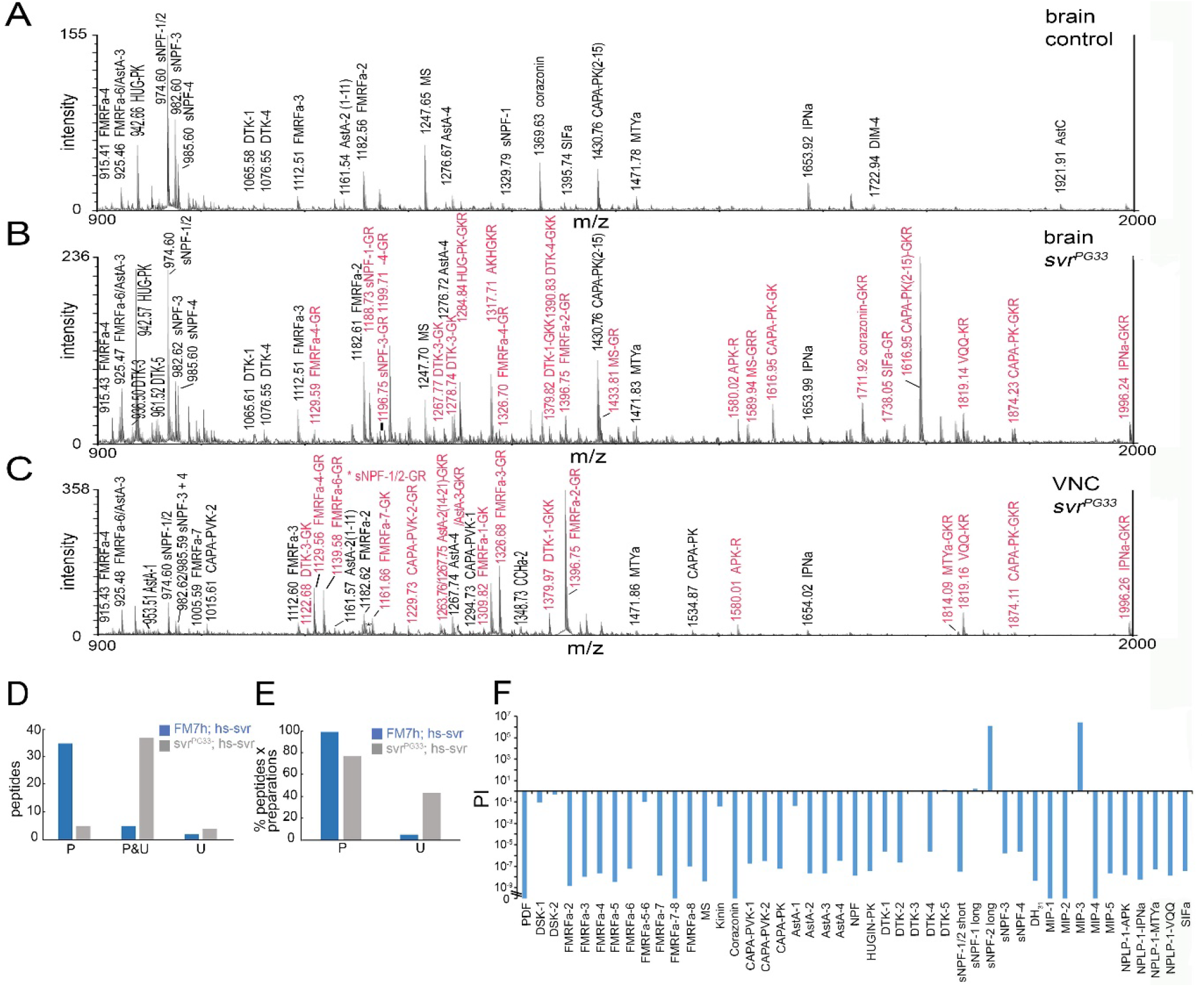
Comparative Peptidomics betwenn svr mutant and control flies. **A-C)** MALDI-TOF spectra from direct peptide profiling. **A)** Example for central brain tissue of a FM7h; hs-svr control fly in the mass range of 900-2000 Da. Note that all detected neuropeptides are fully processed, indicated by black letters. **B)** Example for central brain tissue of a svr^PG33^;hs-svr mutant fly. Note that the profile for fully processed neuropeptides (black letters) is very similar to that for controls in A). Yet, in addition, peaks indicating C-terminally extended forms (red letters) are present. **C)** Similar C-terminally extended forms are also visible in other tissues, here a ventral nerve cord of a svr^PG33^; hs-svr mutant fly although a partly different neuropeptide complement is present. **D-E)** Summary of the processed and unprocessed neuropeptide forms identified by direct peptide profiling. **D)** Number of different peptides found only in either a fully processed (left) or fully unprocessed C-terminally extend form (right), or in both forms (middle) throughout all analysed samples. E) Same data as in D), but now presented as fractions per individual preparations. The data show that unprocessed peptides are more abundant in svr^PG33^; hs-svr mutants than in FM7h;hs-svr control flies, for which unprocessed peptides occurred only for a few specific peptides (see text and SFig. 1). In contrast, processed peptides, processed peptides are more abundant in FM7h; hs-svr control flies than in svr^PG33^;hs-svr mutants. **F)** Processing index (PI) for the neuropeptides identified by LC-MS in mutant svr^PG33^;hs-svr and control FM7h/hs-svr flies. A PI>1 indicates that more processed peptide is present in the mutant than in the control. A PI<1 indicates that more unprocessed peptide is present in the mutant than in the control. PI=0 indicates that only unprocessed peptide was present in the mutant.

In total, 43 different peptides (equalling about 80% of the mass-spectrometrically confirmed *Drosophila* neuropeptides (6)) were identified by mass match (± 0.5 Da). In *FM7h*; *hs-sv*r control flies, 35 of these peptides were only found in their processed bioactive form (Fig. 2D-E, Supporting Fig. 1). For five peptides (CAPA-PVK-2, CCHa-2, FMRFa-3, FMRFa-4 and kinin) both processed bioactive and C-terminal extended forms were found. For AstA-2 and CCAP, only mass peaks matching the unprocessed peptide carrying a C-terminal GKR and GRKR extension were identified. Due to the prevalence of nonpolar amino acids in AstA-2 (LPVYNFGLamide) and CCAP (PFCNAFTGCamide, carrying an internal cystein bridge), both peptides are notoriously difficult to detect by direct MALDI-TOF peptide profiling without prior chemical modification. Yet, the C-terminal Arg of the unprocessed peptide likely increased ionisation efficiency during the MALDI process (35). In 99.0% of the detected peptide peaks over all preparation, fully processed forms were found, while peptides carrying a C-terminal basic amino acid extension occurred in only 5% of the peptides*preparations (Fig. 2D-E, Supporting Fig. 1).

In *svr*^*PG33*^, *hs-svr* experimental flies, 37 neuropeptides were detected by mass match in both processed bioactive and C-terminal extended form, of which APK-R, AstA-4, CAPA-PK^(2-15)^-GKR, CAPA-PVK-1, corazonin-GKR, FMRFa-2-GR, FMRFa-3-GR, FMRFa-4-GR, FMRFa-6, FMRFa-6-GR, HUG-PK-GKR, IPNa-GKR, myosuppressin-GRR, SIFa, sNPF-1^(4-11)^-GR were sequence-confirmed by MS/MS fragmentation. Only five neuropeptides (Ast-C, CCHa-2, FMRFa-8, MIP-3 and MIP-5) were exclusively found in their processed bioactive form. Four neuropeptides (CCAP, kinin, MIP-2, PDF) were exclusively found in their unprocessed C-terminally extended form (Supporting Fig. 1). In 76.8% of the peptides*preparations, fully processed forms were found, while peptides carrying a C-terminal basic amino acid extension were found in 43.2% of the peptides*preparations (Fig. 2D-E, Supporting Fig. 1).

The qualitative results obtained by direct peptide profiling indicate that the occurrence of unprocessed C-terminally extended neuropeptides is highly increased across the peptidome in *svr*^*PG33*^, *hs-svr* flies, suggesting that dCPD is involved in the processing of most if not all neuropeptides. To obtain a more quantitative measure of the effect of a dCPD loss-of-function, we next used nanoLC-ESI-MS/MS and measured relative differences in the detection levels of processed and unprocessed peptides in brain extracts of *svr*^*PG33*^, *hs-svr* mutant flies and *FM7h*; *hs-sv*r controls. Representative raw data are shown for allatostatin A and FMRF-like peptides in Supporting Fig. 2. For each neuropeptide, we calculated a processing index (PI) as a measure of the relative differences of processed vs. unprocessed peptides between *svr*^*PG33*^, *hs-svr* mutant flies and *FM7h*; *hs-sv*r controls. The results are summarised in Fig. 2F, the ratios (R) for the peak areas between unprocessed and processed peptides are shown in Supporting Fig. 3. With exception of MIP-3 and the long forms of sNPF-1 and -2, all peptides had a PI smaller than 0.1, indicating that dCPD is required for C-terminal trimming and peptide processing. In *FM7h*; *hs-sv*r controls, all peptides were exclusively detected in their fully processed form, with exception for AstA-1 and NPLP1-IPNa. Unprocessed AstA-1 was found in one out of three samples (R = 12.5) and unprocessed NPLP-1-IPNa occured in two out of three samples (R=12.4 and 1.9). In *svr*^*PG33*^, *hs-svr* mutant flies, most peptides were found in both processed and unprocessed form, confirming the results from MALDI-TOF peptide profiling. For the mutant, LC-MS also confirmed the MALDI-TOF detection of exclusively unprocessed kinin, MIP-2, and PDF. Also corazonin, MIP-1 and MIP-4 were exclusively detectable in their unprocessed form. Further, MIP-3 but not MIP-5 and FMRFa-8 was found in exclusively processed form in the mutant. Like MIP-3, also the long forms of sNPF-1 (sNPF-1_1-11_ AQRSPSLRLRFa) and sNPF-2 (sNPF-2_1-19_ WFGDVNQKPIRSPSLRLRFa) were only detectable in the mutant, but occured with a smaller peak area also in the unprocessed forms. This explains the positive PIs for these peptides (Fig. 2F). The presence of only processed MIP-3 in mutant flies is difficult to account for. sNPF-1_1-11_ as well as sNPF-2_1-10_ has been found at lower signal intensity in previous peptidomic studies on larval or adult *Drosophila* CNS (32, 33, 36–39). In contrast, sNPF-2_1-19_ had not been biochemically identified so far, although this peptide was early on predicted from the *Drosophila* genome (40). Thus, the presence of processed and unprocessed sNPF-1_1-11_ and -2_1-19_ suggests that prior efficient dCPD action is required for a proper intrapeptide processing at the monobasic arginine to create the sequence-identical short sNPF forms (sNPF_4-11_, sNPF-2_12-19_ SPSLRLRFa).

**Figure 3:**
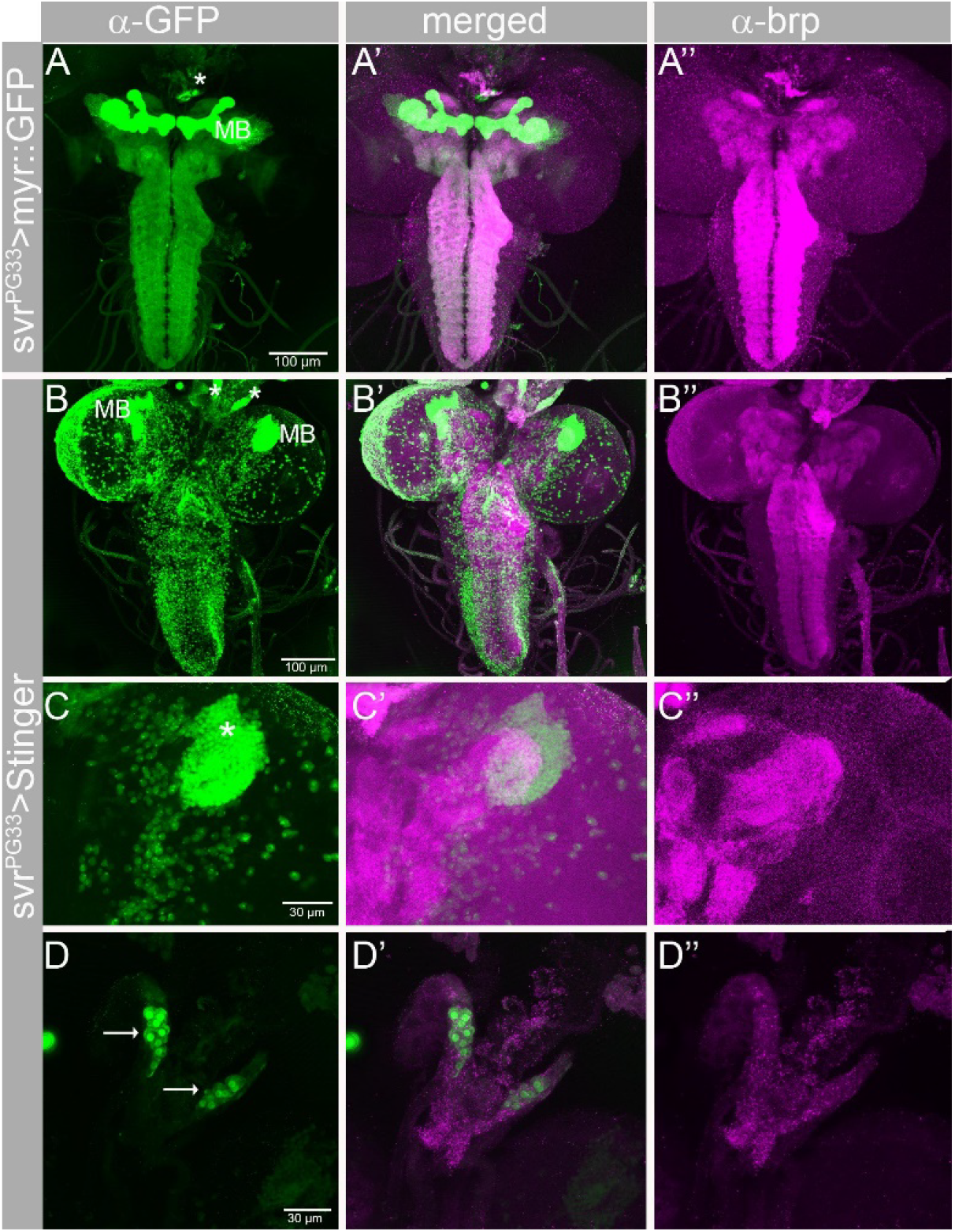
svr expression pattern based on the intragenic PG33 insert. The Gal4 sequence contained within the PG33 GawB insert was used to drive either a membrane-bound (myr::GFP, A) or nuclear (Stinger) form of GFP (B-D) in the larval CNS. Staining with mAb nc82 against the synaptic protein bruchpilot was used to mark neuropil areas (A’’-D’’). Maximum projections of confocal stacks. **A)** The whole neuropile is uniformly innervated by svr^PG33^-expressing neurons, only the densly packed mushroom bodies (MB) stick out. In the attached ring gland, the endocrine cells producing adipokinetic hormone are also strongly labelled (asterisk). **B)** The nuclear staining shows that many but by far not all neurons express svr^PG33^. Again, the Kenyon cells forming the mushroom bodies (MB) as well as the AKH cells (asterisks) are conspicuous. **C)** Close up of the MB shown in B) shows the densly packed Kenyon cell somata (asterisk) in the dorsal protocerebrum. **D)** Close up of the AKH cells in B), located in the corpora cardiaca portion of the ring gland.

### *svr* is broadly expressed throughout the larval and adult CNS

The peptidome-wide impairment of neuropeptide processing in *svr*^*PG33*^, *hs-svr* mutants suggest that dCPD is broadely expressed in peptidergic neurons throughout the CNS. To test this, we took advantage of the intragenic Gal4 sequence contained within the inserted PGawB element of *svr*^*PG33*^(25), and drove the expression of either a membrane-bound or nuclear form of UAS-GFP. In both the larval and adult CNS, a large number of neurons innervating basically all neuropiles expressed *svr*^*PG33*^-Gal4-driven GFP (Fig 3A). This is in line with the idea that many *Drosophila* neurons express peptide co-transmitters (41). Due to the broad expression, it was impossible to identify specific arborisation patterns with exception of the tightly packed Kenyon cells with their parallel projections in the mushroom bodies (Fig. 3A-C). These neurons are known to express sNPF (36, 42). As sNPF processing was impaired in *svr*^*PG33*^, *hs-svr* (Fig. 2F, Supporting Fig. 1), dCPD likely plays a role in peptide processing in these higher order integration centers. Outside of the CNS, the endocrine AKH cells in the glandular part of the corpora cardiaca, the proximal part of the larval ring gland were strongly labelled (Fig. 3A, D). This is compatible with earlier genetic manipulations of dCPD levels in these cells, which significantly affected AKH processing (24).

### dCPD is required for starvation-induced hyperactivity

The expression pattern of *svr*^PG33^ suggested that dCPD is expressed in AKH-producing endocrine cells of the corpora cardiaca. AKH has metabolic functions and is required for starvation-induced hyperactivity (43, 44). To address the functional requirement of dCPD in AKH processing, we tested whether starvation-induced hyperactivity is affected in *svr*^*PG33*^, *hs-svr* mutant flies (Fig. 4A-B). At ZT 12 (lights off), flies were placed into monitor tubes containing either only agarose (starvation condition) or agarose and sucrose (control condition). All genotypes showed significant starvation-induced hyperactivity during the experiment on agarose (Fig. 4A). *svr*^*PG33*^, *hs-svr* mutant flies increased their locomotor activity already during the first day (12-24h of starvation), while controls still showed normal day activity. During the following night (24-36h of starvation), also control flies increased their locomotor activity to a significant level (Fig. 4B). The maximal levels of activity of *svr*^*PG33*^, *hs-svr* mutant flies were quite stable throughout the 72h under starved conditions, and locomotor activity still showed daily rhythmicity with distinguishable peaks during morning and evening and intermediate siesta phase from 24-60h. In contrast, locomotor activity of control flies peaked during the second night after 24-36h of starvation, and then steadily declined until all flies had died at 72h without any sign of a siesta phase. In other words, *svr*^*PG33*^, *hs-svr* mutant flies showed reduced hyperactivity compared to controls, in which starvation-induced hyperactivity completely overruled the daily activity pattern. This phenotype is compatible with the assumption of a hypomorphic alteration of AKH processing.

**Figure 4:**
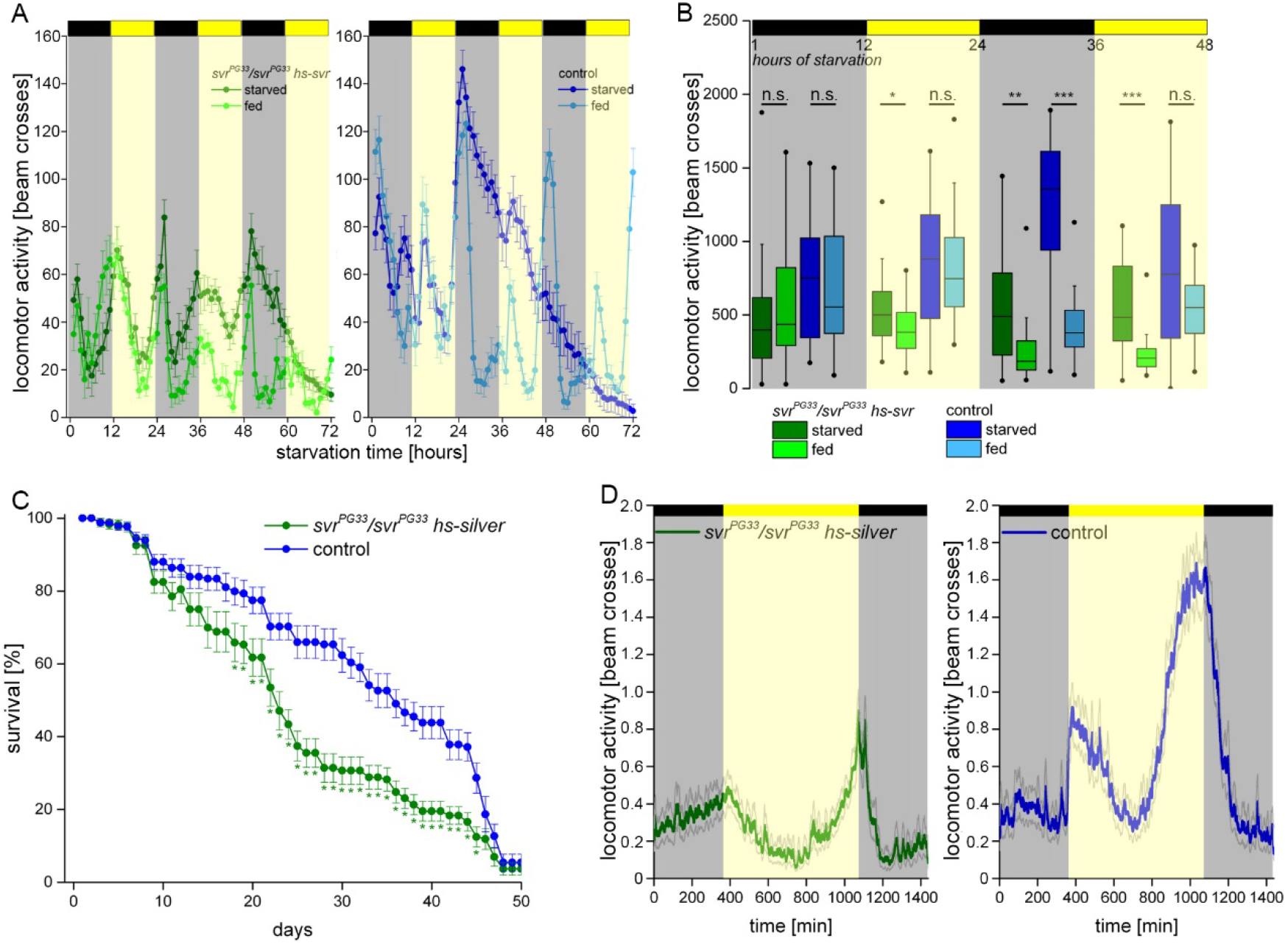
dCPD is required for starvation-induced hyperactivity. **A)** svr^PG33^, hs-svr mutant flies showed an early onset of starvation-induced hyperactivity (∼20h) during the first day without food. In contrast, control flies started starvation-induced hyperactivity later, during the second night after ∼30h of starvation. **B)** Statistic comparison of averaged locomotor activity for the first two nights and days after starvation onset. n=23-29. **C)** Survival is significantly lower between day 18 and day 45 in svr^PG33^, hs-svr mutant flies compared to control flies. **D)** Locomotor activity in svr^PG33^, hs-svr mutant flies is significantly reduced compared to control flies, especially during morning and evening activity bouts. Yet, daily rhythmicity remains unaffected. n=53-61.

### Loss of dCPD affects survival rate and general activity levels

To test whether *svr*^*PG33*^, *hs-svr* mutant flies show an impaired general health state, we analysed their locomotor activity level and life span. Both *svr*^*PG33*^, *hs-svr* mutant and control flies showed a maximal life span of around 50 days under *ad libitum* feeding conditions at 25°C. However, *svr*^*PG33*^, *hs-svr* mutant flies showed a significantly lower survival rate between day 18 and day 45 compared to controls (Fig. 4C). Next, we analyzed locomotor activity and rhythmicity. Control flies displayed the characteristic bimodal activity pattern with increased morning and evening activity and a siesta phase during mid-day. *svr*^*PG33*^, *hs-svr* mutant flies showed the same rhythmicity, yet an overall reduced activity pattern, with dampened morning and evening activity bouts (Fig. 4D).

### dCPD is required for appetitive but not aversive gustatory preference

Neuropeptides act both as signalling or modulatory substances within chemosensory input pathways in *Drosophila* (45–48). We therefore used larval preference assays (49) to functionally test whether dCPD plays a role in processing peptides involved in gustatory or olfactory signalling. First, we tested whether the lack of dCPD affects the preference for fructose, a sugar that provides nutritional value and sweetness (50). Individual larvae were analysed over time using the FIM tracking system (51). *svr*^*PG33*^, *hs-svr* mutant larvae showed approach behavior for fructose, however, the performance was reduced compared to control larvae in the course of time (Fig. 5A) and significantly reduced after 5min (Fig. 5D). To test whether a lack of dCPD generally affects gustatory responses, we challenged larvae with high salt concentrations. As expected, control larvae significantly avoided high salt concentrations. Similarly, *svr*^*PG33*^, *hs-svr* mutant larvae showed avoidance behavior suggesting that the lack of dCPD affected the larval response specifically to sugar or appetitive substances instead and not taste responses in general (Fig.5B, E). Next, we tested larvae for their innate olfaction. *svr*^*PG33*^, *hs-svr* mutant larvae and control larvae performed indistinguishable over the course of time and approached amylacetate (Fig. 5C, F). Our results suggest that hypomorphic dCPD levels do not affect chemosensory sensory processing in general, but rather gustatory responses to sugar.

**Figure 5:**
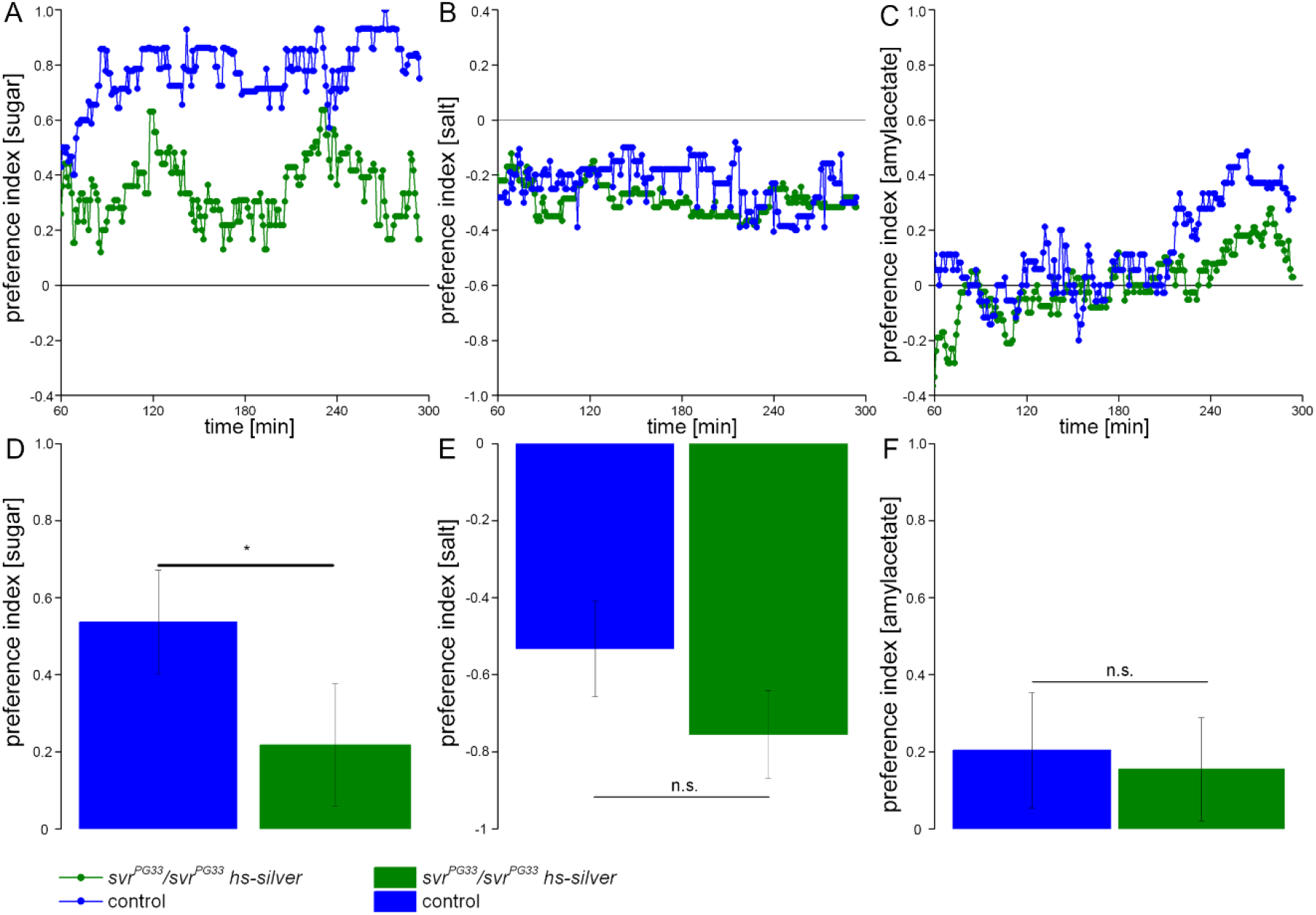
dCPD affects appetitive, but not aversive taste or odor responses. **A,D)** svr^PG33^, hs-svr mutant larvae showed a reduced preference behavior for fructose compared to control larvae. (B,E) svr^PG33^, hs-svr mutant larvae avoided high salt concentration similar to control larvae over the course of time. (C,E) Odor response to amylacetate was indistinguishable between svr^PG33^, hs-svr mutant larvae and control larvae. n=30-42.

### Evidence for an independent loss of CPE but not CPD in holometabolous insects

CPE, the major neuropeptide processing CP in mammals, has also been found in molluscs (52) and nematodes (53, 54), but appears to be absent in *Drosophila* (22). To test whether the lack of CPE applies to insects in general, we performed a non-exhaustive insect genome BLAST search based on CPE sequences from mouse and *C. elegans* (EGL-21), and CPD sequences from *Drosophila*. Obtained insect CPE sequences were then used to refine the search. In total, we obtained 310 predicted full CP sequences from insects covering all major insect orders. We then calculated a maximum likelihood tree, which clustered CPE and CPD/CPM separately of each other (Supporting Fig. 4). CPDs have three CP domains, and are therefore significantly larger than CPEs and CPMs. This criterion was used to separate CPDs from CPMs which otherwise show high sequence similarity between their active CP domains. Our analysis certainly does not include all available sequence information for insects and we did not specifically search for CP sequences within individual genomes. Our BLAST search yield CPD sequences for most insect orders (Table 1), except for diplurans, mayflies (Ephemeroptera), dragon flies (Odonata) and stick insects (Phasmatodea) for which little genomic data is available and CPD sequences are likely to be found by a more thorough analysis. A conservative interpretation of the data is that all insect orders for which abundant genomic data is available possess CPD (Table 1). CPE was found in the basal Entognatha and in the majority of insect orders including most if not all hemimetabolous orders as well as the holometabolous beetles and the derived Lepidoptera (butterflies, moths and allies) (Table 1). Interestingly, putative CPE sequences could not be identified from the two large holometabolous orders which contain the majority of insects with sequenced genomes: the Diptera (mosquitoes, flies and allies) and the Hymenoptera (bees, ants, wasps and allies) (Table 1). In other words, CPE seems to be common throughout the insects, but seems to have been independently lost at least two times: in the basal holometabolous Hymenoptera, as well as the derived holometabolous Diptera. A puzzling finding not concurrent with this notion is the presence of putative CPE sequences in flies belonging to the *Tephritidae* (true fruit flies, though belonging to the acalyptrate group of flies they are not directly related to the *Drosophilidae* (vinegar flies)). The size and sequence classifies these predicted CPs as CPEs. Furthermore, we also found a true CPD gene for the respective tephritid species (snowberry fruit fly *Rhagoletis zephyria*, solanaceous fruit fly *Bactrocera latifrons*) speaking against a gene prediction or sequencing error. However, sequence similarities between the CP domains in CPE and CPD including the catalytic triad and Zn^2+^ binding site are high and CPD-encoding genes are long and contain numerous interons (22). It is clear that further studies are required to confirm the surprising putative presence of CPE in tephritids.

**Table 1:**
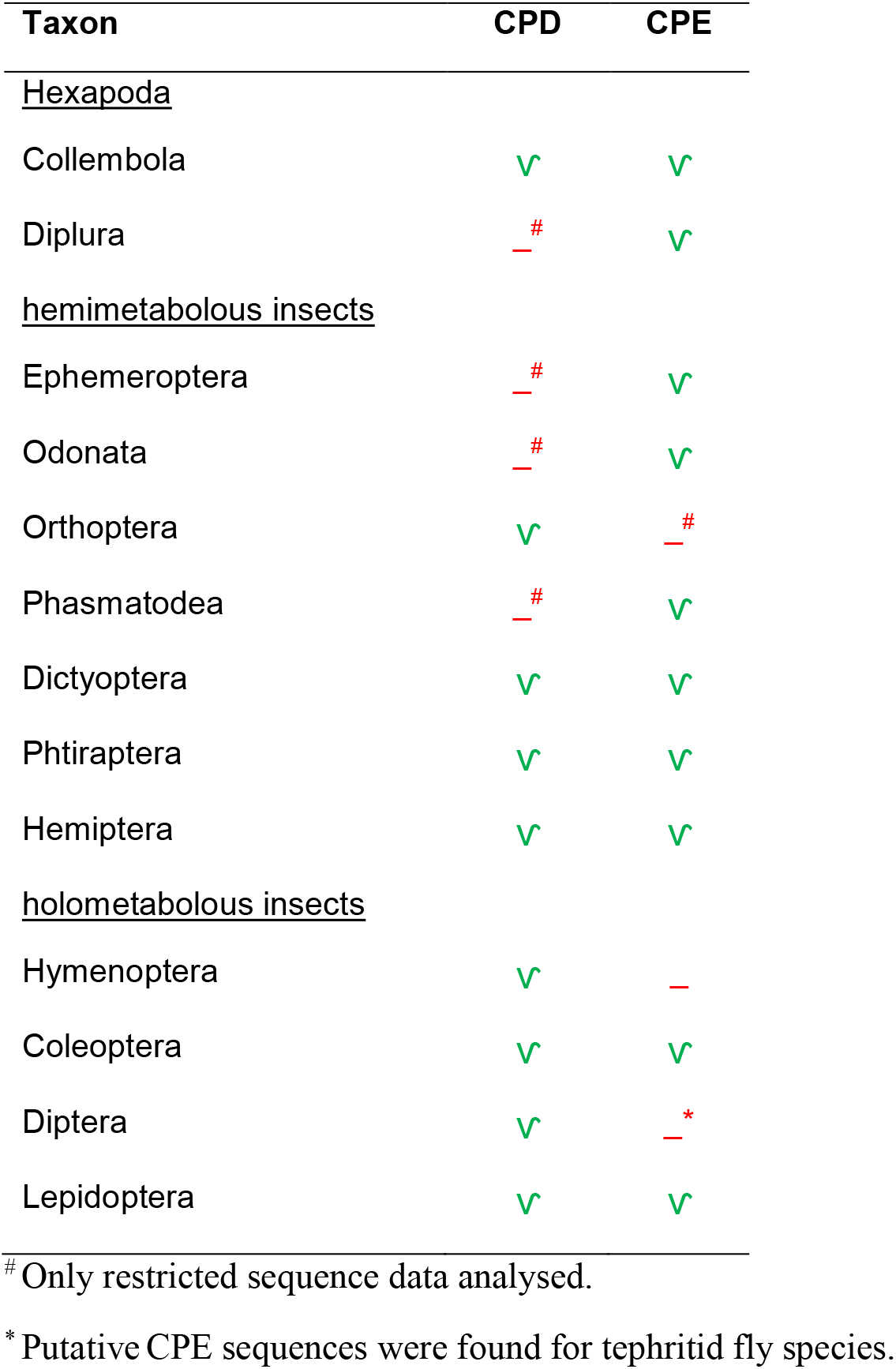
Taxonomic distribution of identified putative insect CPD/CPE sequences.

| **Taxon** | **CPD** | **CPE** |
| --- | --- | --- |
| <u>Hexapoda</u> |  |  |
| Collembola | ✓ | ✓ |
| Diplura | —<sup>#</sup> | ✓ |
| <u>hemimetabolous insects</u> |  |  |
| Ephemeroptera | —<sup>#</sup> | ✓ |
| Odonata | —<sup>#</sup> | ✓ |
| Orthoptera | ✓ | —<sup>#</sup> |
| Phasmatodea | —<sup>#</sup> | ✓ |
| Dictyoptera | ✓ | ✓ |
| Phthiraptera | ✓ | ✓ |
| Hemiptera | ✓ | ✓ |
| <u>holometabolous insects</u> |  |  |
| Hymenoptera | ✓ | — |
| Coleoptera | ✓ | ✓ |
| Diptera | ✓ | —<sup>*</sup> |
| Lepidoptera | ✓ | ✓ |

<sup>#</sup> Only restricted sequence data analysed.
<sup>\*</sup> Putative CPE sequences were found for tephritid fly species.

## Discussion

N/E metallocarboxypeptidases of the M14 family catalyse the second step in prepropeptide processing by removing the C-terminal mono-or dibasic cleavage signal that remains after proprotein convertase cleavage of the propeptide (16). Peptidomic studies detected a large and broad accumulation of C-terminally extendend neuropeptides in *Cpe*^*fat*^*/Cpe*^*fat*^ mice (13, 55, 56) and *egl-21* mutant *C. elegans* (54). This indicated that CPE is the key carboxypeptidase for neuropeptide processing in these phylogenetically very distant animals. A gene coding for CPE could however not been found in the fruit fly *Drosophila* (22, 57), the only insect in which neuropeptide processing is studied in more detail (6). Several lines of evidence indicated that instead dCPD encoded by *svr* has taken over the key function of neuropeptide processing in *Drosophila*. First, the second domain of dCPD has enzymatic properties similar to those of mammalian CPE (22). Next, ectopic expression of *svr* in the endocrine AKH-producing cells reduced the levels of naturally occuring C-terminally extended AKH (24). Lastly, flies with disrupted *svr* gene are embryonically lethal (21, 23) and show deficits in various neuropeptide-regulated behaviours (23). However, direct evidence for an involvement of CPD in neuropeptide processing was so far lacking for any species, including the fruit fly. Hence, a main result of our peptidomic study is the demonstration of the ability of dCPD to efficiently process a broad variety of neuropeptides. This raises the question whether the failure of mouse CPD to fully rescue a deficiency of CPE (17) is caused by an incomplete overlap of the CPD and CPE expression patterns in peptidergic neurons rather than by a reduced efficiency of mouse CPD to process neuropeptides.

The early lethality of *svr* mutants (21, 23) suggests that dCPD function during development cannot be substituted by dCPM. Our finding that *svr*^*PG33*^ mutant larvae and pupae required daily heat-shock rescue to develop into adults is in line with the importante of *svr* during development. Yet, once adult, *svr*^*PG33*^ mutants survived without heat-shock even though the survival rate was significantly reduced. Our RT-PCR results suggest that this reduced viability is due to a hypomorphic background expression of the *hs-svr* rescue construct even at lower temperature, and not due to a loss of dCPD requirement in adult flies. We had previously observed a similar phenomenon for *amon* (dPC2) deficient flies rescued by a *hs-amon* construct (9). We hypothesise that the production rate of neuropeptides is significantly lower in adult flies than in the very fast growing larva or developing pupa, in which the background expression of either *hs-amon* or *hs-svr* may not be sufficient to provide the required increasing supply in neuropeptides during development. Taken together, our results suggest that dCPD is a key if not the sole CP involved in neuropeptide processing in *Drosophila*.

Given that most animals have additional genes for PC1/3 and CPE, it is surprising why apparently *Drosophila* during evolution has kept only one gene for a proprotein convertase (dPC2, *amon*, (9) and one for a CP (*svr*), considering that a mutation in either gene is lethal. One reason for keeping CPD and not CPE could be the potential higher versatility of CPD with its two bioactive domains and membrane-anchored and free isoforms which have unique functions, substrate specificities and specific pH optima (18, 24). It is tempting to speculate that dCPD has further functions additional to neuropeptide processing, which cannot be executed by CPE with its single bioactive domain. This is compatible to the finding that hypomorphs have phenotypes in wing morphology (23) and biogenic amine pools (26). This hypothesis is also supported by our finding that obviously CPE has been independently lost at least twice during insect evolution in Hymenoptera and Diptera, while CPD seems to be present in all insect taxa.

Based on the expression pattern of the *svr*^*PG33*^ transgene, dCPD is broadly expressed in the CNS and neuroendocrine organs. The expression in the Kenyon cells of the mushroom body, an important brain center for learning and memory, is in line with impaired long-term memory in a courtship assay (24) and olfactory memory formation (28). Similarly, knock-out of CPE in mice impairs olfactory social learning, object recognition memory and performance in the Morris water maze (58). The expression of *svr*^*PG33*^ in the endocrine AKH cells as well as its effect on AKH-mediated starvation-induced hyperactivity, indicates a role of of dCPD in AKH processing after dPC2 cleavage (10). This is in line with the effect of ectopically expressed dCPD on AKH processing as shown by direct peptide profiling (24).

Impairment in insulin signalling increases life span (59, 60), and dCPD has been implicated in insulin processing (28). Yet, we found the maximum life span of the *svr*^*PG33*^ hypomorphs is simlar and survival rate is decreased. This suggests a further, unidentified role of dCPD outside of peptide processing or an involvement of other yet unknown neuropeptides in the regulation of life-span. These neuropeptides could be identified by cell-specific *svr* knock-out via CRISPR-Cas9 (61) or knock-down via RNAi, as the *svr*^*PG33*^ driver line lends itself as a tool to generally impair peptide signalling in neurons even if these express multiple co-expressed or unknown peptides.

Taken together, our results show that dCPD is a key enzyme for neuropeptide processing in *Drosophila*, and is required for proper peptide-regulated behaviour. dCPD also may have functions unrelated to neuropeptide processing which led to an evolutionary retention of CPD in insect genomes.

## Experimental procedures

### Fly strains

The *svr* null mutant *y*^*1*^ *w*^*\**^ P{*w*^*+mW*^ ^*hs*^=*GawB*}*svr*^*PG33*^/FM7h (25) was a kind gift of Galina Sidyelyeva and Lloyd Fricker. *w*^*1118*^ controls, *10xUAS-IVS-myr::GFP, UAS-Stinger* were obtained from Bloomington Stock Center.

Flies were kept on standard food at either 18°C or 25°C and an LD12:12 light cycle and a relative humidity of 60%.

### Generation of hs-svr flies

To rescue the lethal *svr*^PG33^ mutant phenotype, we generated flies carrying a *hs-svr*^1A-2-3-t2^ genomic insert. The pUAST-*svr* (1A-2-3-t2) plasmid ((24), kind gift of Galina Sidyelyeva and Lloyd Fricker) was digested using *EcoRI* and *XbaI*, resulting in two fragments consisting of the 5’-2174 bp and the 3’-2351 bp part of the svr (1A-2-3-t2) construct. Then, the pCaSpeR-hs vector (62) was digested using *EcoRI* and *XbaI*, and the 3’-2351 bp part was inserted. After ligation, the resulting plasmid was digested with *EcoRI*, and the 5’-2174 bp fragment was inserted and ligated. The correct orientation of the fragments and sequence was confirmed by Sanger sequencing, and the resulting *hs-svr*^1A-2-3-t2^ construct was introduced into the germline of w^1118^ flies by BestGene (Chino Hills, CA, USA) using standard P-element transformation. Several independent transformant lines were obtained which were used to generate *w*^*1118*^*; hs-svr*^*1A-2-3-t2*^ and *y*^*1*^ *w*^*\**^ P{*w*^*+mW*^ ^*hs*^=*GawB*}*svr*^*PG33*^/FM7h; *hs-svr*^1A-2-3-t2^ lines.

### Heat-shock rescue

Eggs of *y*^*1*^ *w*^*\**^ P{*w*^*+mW*^ ^*hs*^=*GawB*}*svr*^*PG33*^/FM7h; *hs-svr*^1A-2-3-t2^ flies were collected every morning, kept at room temperature and heat-shocked daily for 30 min at 37°C in a water bath. Larvae for the behavioural assays were obtained by heat-shocking for four days, followed by two days at 18°C to minimise background expression of *hs-svr*. Homozygous mutant larvae were identified by the light mouth parts and unpigmented denticle bands due to the *y*^*1*^ allele, while larvae carrying FM7h showed stronger pigmentation due to the *y*^*31*^ allele of FM7h (63). To obtain adults, heat-shocking was continued until eclosion. After eclosion, flies were kept at 18°C, until male nervous tissue was dissected and processed as described. For behavioural and life span assays, flies 4-7 respectively 1 day after eclosion were used. FM7h controls were distinguished from *svr*^*PG33*^ mutants by the presence of barred (*B)* and orange-coloured eyes.

### RT-PCR

To test for background expression of *hs-svr*, total RNA was extracted from heads of five adult males per genotype using the Quick-RNA MicroPrep Kit (Zymo Research, Irvine, USA) according to manufacturer’s instructions. Heads were cut off, collected in a microtube containing 300 µl RNA lysis buffer on ice, and homogenised with a plastic pestle. Total RNA was eluted in 8µl RNAse-free water. For cDNA synthesis, the QuantiTect Reverse Transcription Kit from Qiagen (Venlo, Netherlands) was used. All steps were performed following the manufacturer’s protocol. Genomic DNA was removed by adding 1 µl of gDNA wipeout buffer to 6 µl of the eluted RNA. Following incubation at 42 °C for 2min, the samples were placed for 2min at 4 °C and 3 µl of a mastermix composed of 8 µl RT Buffer, 2 µl RT Primer Mix and 2 µl reverse transcriptase was added. Reverse transcription was performed for 30 min at 42 °C, followed by 3 min at 95 °C and 2min at 4 °C. Finally, 40 µl water were added and cDNA samples were stored at −20 °C.

cDNA was PCR-amplified using a JumpStart REDTaq ReadyMix (Sigma-Aldrich) and *svr-, GawB*- and *Gaw-svr*-specific primers (see Supplemental table 1). *α-tubulin* was used as internal control. The PCR program consisted of 5min at 95°C, followed by 30 cycles of 30 s at 95°C, 30 s at 60°C, 60 s at 72°C, followed by a final extension for 5 min at 72°C.

### Direct peptide profiling via MALDI-TOF mass spectrometry

Direct peptide profiling was carried out according to our standard protocol (64). In brief, the brain and ventral ganglion (VG) of adult male flies 14 days after eclosion and last heat-shock was dissected in HL3.1 saline: 170 mM NaCl, 5 mM KCl, 1.5 mM CaCl_2_, 4 mM MgCl_2_, 10 mM NaHCO_3_, 5 mM trehalose, 115 mM sucrose, 5 mM HEPES, pH 7.5 (65). Using microscissors, the brain was further divided into optic lobes or central brain. Tissues were then transferred by pulled glass capillaries to a stainless steel MALDI target, remaining saline was removed and the tissues were let to dry. In case of excess salt deposits, a small droplet of ice-cold water was added to the dried tissues and removed after about 1s to desalt the sample.

Then, 200 nl matrix (saturated solutions of recrystallized a-cyano-4-hydroxycinnamic acid (CHCA) in 30% MeOH: 30% EtOH and 40% water (v:v:v)) was added per tissue and let to dry. MALDI-TOF mass spectra were acquired in positive ion mode on an Applied Biosystems 4800 Plus MALDI-TOF/TOF mass spectrometer (Applied Biosystems/MDS Sciex)). Samples were analyzed in positive reflector mode within a mass window of 850-3000 Da. Some mass peaks were additionally fragmented in MS/MS mode using PSD. Laser power was adjusted manually to provide optimal signal-to-noise ratio. Raw data were analyzed using Data Explorer 4.10 software (Applied Biosystems/MDS Sciex).

### Sample preparation for NanoLC-ESI-MS/MS

After eclosion, flies were kept for 5 days without heat-shock. Then, brains, ventral nerve cords and guts of males were dissected in HL3.1 saline on ice and transferred by a needle to a frozen microtube in a laptop cooler. Per sample, 30 brains plus ventral nerve cords or 30 guts were pooled and stored at - 80°C until extraction. For extraction, 50 µl of a methanol, water, trifluoracetic acid mixture (90/9/1 v/v/v) were added to the frozen tubes, followed by 3 min in an ice-cold ultrasonic bath and 30 min incubation on ice. Afterwards, the samples were centrifuged for 15 min at 15.000 g (Hettich Mikro 200, Tuttlingen, Germany), and the supernatant was transferred to a new microtube and vacuum-dried (Uniequip Univapo 100H, Planegg, Germany).

The extracts were prepurified on self-made StageTips (66) using 3M Empore C18 material (ChromTech Inc., Apple Valley, MN, USA) and 200 µl pipet tips. The columns were activated with 50 µl of 100% acetonitrile, and equilibrated with 50 µl of 10 mM HCl. The extracted samples were dissolved in 30 µl of 10 mM HCl, sonicated in an ultrasound bath and applied onto the equilibrated StageTips. After washing with 50 µl 10 mM HCl, peptides were eluted with 30% acetonitrile and dried in a vacuum concentrator (SpeedVac, Eppendorf). Low-binding plastic was used throughout.

### NanoLC-MS/MS Analysis

Peptides were dissolved in 2 % acetonitrile, 0.1 % formic acid. NanoLC-MS/MS analyses were performed on an Orbitrap Fusion (Thermo Scientific) equipped with a PicoView Ion Source (New Objective) and coupled to an EASY-nLC 1000 (Thermo Scientific). Peptides were loaded on capillary columns (PicoFrit, 30 cm x 150 µm ID, New Objective) self-packed with ReproSil-Pur 120 C18-AQ, 1.9 µm (Dr. Maisch) and separated with a 30-minute linear gradient from 3% to 40% acetonitrile and 0.1% formic acid and a flow rate of 500 nl/min.

Both MS and MS/MS scans were acquired in the Orbitrap analyzer with a resolution of 60,000 for MS scans and 15,000 for MS/MS scans. A mixed ETD/HCD method was used. HCD fragmentation was applied with 35 % normalized collision energy. For ETD calibrated charge-dependend ETD parameter were applied. A Top Speed data-dependent MS/MS method with a fixed cycle time of 3 seconds was used. Dynamic exclusion was applied with a repeat count of 1 and an exclusion duration of 10 seconds; singly charged precursors were excluded from selection. Minimum signal threshold for precursor selection was set to 50,000. Predictive AGC was used with AGC a target value of 2e5 for MS scans and 5e4 for MS/MS scans. EASY-IC was used for internal calibration.

In total, three (controls) and two (mutants) biological samples were measured in technical duplicates. All chemicals used were of HPLC grade; low-binding plastic was used throughout.

### Data Analysis

Data analysis was performed with PEAKS Studio 8.5 (Bioinformatics Solutions Inc. (67)). Parent mass tolerance was set to 8 ppm, and fragment mass tolerance to 0.02 Da. Pyro-Glu (N-term Q), oxidation (Met), carbamidomethylation (Cys) and amidation (C-term) were allowed as variable modification. A maximum number of 5 modifications per peptides were allowed. Searches were performed against a custom neuropeptide database that contained all known *Drosophila* and suggested prepropeptides, processing enzymes as well as neuropeptide receptors. Results were filtered to 1% PSM-FDR. To calculate the processing index (PI), we first calculated the mean ratio R of the peak area (PA) between the fully processed and unprocessed form (PA_fully processed_/PA_C-terminally extended_) for the mutant and control samples respectively. These ratios were then used to calculate the PI for each peptide (PI = R_*svrPG33* mutant_/R_*FM7h* control_) to correct for possible differences in the ionisation probabilities between processed and unprocessed peptides.

### Immunostainings

*svr* ^*PG33*^/FM7h flies were crossed with w*,10x*UAS-IVS-myrGFP* or *w*;UAS-Stinger*. Larval CNS were dissected in HL3.1 saline (65) and fixed for 2h in 4% paraformaldehyde in 0.1M PBS at room temperature. Then, brains were washed in 0.1 M PBS with 0.3% TritonX (PBT), and incubated in PBT+5% normal goat serum for 1h at room temperature. Afterwards, primary antibodies (anti-GFP rabbit polyclonal (1:1000, Invitrogen) and anti-brp (nc82) mouse monoclonal (mAb) (1:100)) in PBT+5% NGS were added overnight at 4°C on a shaker. Preparations were then washed 5 x for at least 1h in 1x PBS at room temperature. Fluorophore-coupled secondary antisera (goat-anti-rabbit-Alexa488 and goat-anti-mouse-Dylight 649, Dianova GmbH, Göttingen Germany), diluted 1:1000 in PBT+5% NGS) were added and preparations were incubated overnight at 4°C on a shaker. Next, preparations were washed as above, followed by a final wash in 0.1 PBS and mounting in 80% glycerol in 0.1M PBS. The preparations were analysed with a Leica TCS SPE confocal microscope (Leica SPE, Leica Microsystems Wetzlar, Germany), using ACS APO 20X/0.60 and ACS APO 40X/1.15 objectives. Images were processed with Fiji (68) and Adobe Photoshop CS6 (Adobe Systems, San Jose, California, United States).

### Locomotor activity recording

4-7 day old flies were recorded individually with the *Drosophila* Activity Monitoring (DAM; TriKinetics, Waltham MA, USA) system. To record normal rhythmic behavior, flies were kept in glass tubes with 2% agarose and 4% sugar on one side under standard 12:12 light:dark conditions with light intensities around 100 lux at 20°C and 60% humidity. Fly activity was defined by the number of infrared light beam crosses per minute or per day. Under starved conditions, flies were kept on 2% agarose to avoid dehydration.

### Survival

To test flies for changes in life span, groups of ten flies were kept under normal *ad libitum* standard food conditions at 25°C and 60% humidity and dead flies were counted at the end of each day. Flies were transferred onto fresh food every 3-4 days to circumvent influences due to changes in food quality.

### Larval preference tests

To test larvae for innate odor and taste preference, the FIM (<u>F</u>TIR-based Imaging <u>M</u>ethod) tracking system (51) was used to monitor individual larvae over time. Recordings were made by a monochrome industrial camera (DMK27BUP031) with a Pentax C2514-M objective in combination with a Schneider infrared pass filter, and the IC capture software (www.imagingsource.com). Larval position in relation to the odor or taste stimulus was determined every second. To test larvae for innate odor responses, a thin layer of 1.5% agarose was placed on an acryl plate which was illuminated with infrared light. An odor container (10μl amylacetate) was placed on one side of the agarose layer. A group of five larvae was placed in the neutral mid zone of the agarose layer and larvae were recorded for 5min. To monitor larval responses to fructose or high salt, larvae were placed on a 1.5% agarose layer which contained either 2M fructose or 1.5% sodium chloride on one side.

Preference indices (PIs) were calculated by subtracting the number of larvae on the stimulus-side (ST) from the number of larvae on the no-stimulus side (NO), divided by the total number of larvae: PI=(#ST - #NO) / #TOTAL. Negative PI values indicate avoidance behavior, while positive PI values indicate approach behavior.

### Phylogenomic analysis

A genome-based tblastn search on the NCBI and i5k workspace@NAL web site was performed, using the protein sequences for *Drosophila* CPD and *C. elegans* CPE as a query. The search was then repeated with identified putative insect CPE sequences. Incomplete hits were removed, resulting in a list of 313 CPE/D/M-like sequences that were analysed using MEGA X (69). First, sequences were aligned using the Muscle algorithm, then a maximum likelihood tree employing the JTT matrix-based amino acid substitution model (70) with all sites was calculated with a bootstrap value of 500.

### Statistical analysis

Data were tested for normal distribution using a Shapiro-Wilk test. For the comparison of genotypes, a t-test was used for normally distributed data, and a Wilcoxon Rank Sum test for not-parametrically distributed data. All statistical analyses were done with RStudio, Version 1.0.136 (www.r-project.org). Data plots were made with Origin Pro 2016G, b9.3.226. Significance levels between genotypes shown in the figure refer to raw p-values obtained in the statistical tests.

## Acknowledgments

We thank Lloyd Fricker and Galina Sidyelyeva for the generous gift of plasmids and flies, the Bloomington Stock Center for providing fly lines, and the Drosophila Genomics Resource Center for the generous gift of *svr* constructs, Henry-Marc Bourbon for detailed information on the *svr*^*PG33*^ insert, Lloyd Fricker and Vera Terblanche for helpful discussions, and Susanne Klühspies for lab support. Funded by intramural support of the University of Würzburg.

## Author contributions

DP and CW conceived the project and designed experiments with critical assistance from JTV, AS and JK, YH JTV AS JK and CW performed and analysed the mass spectrometric measurements, YH CW performed immunostainings, DP LT performed and analysed the behavioural assays, TS and GG created the *svr-hs* construct, LT and GG performed RT-PCR, CW performed the phylogenomic analysis, CW and DP wrote the manuscript and generated figure panels, with critical contributions of JTV and AS. All authors commented on the final manuscript version and approved its publication.

## Supporting Information

**Supporting table 1:**
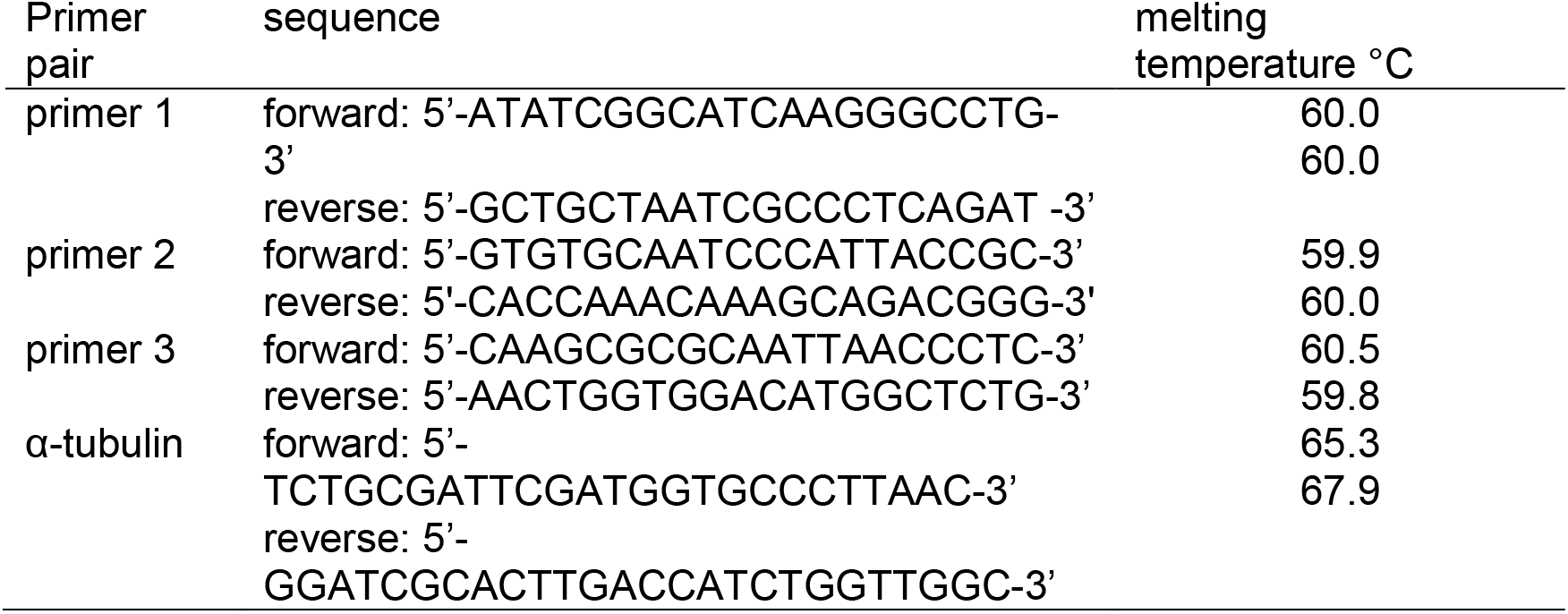
Primer pairs for RT-PCR

**Supporting Figure 1**: Summary of the qualitative MALDI-TOF MS peptide profiling in control (*FM7h; hs-svr*) and mutant (*svr*^*PG33*^*;hs-svr*) flies. The total number of preparations with signals for the respective peptides is given inside the bars, the X-axis shows the relative % of detections. Preparations in which only the respective fully processed peptide was found are shown in gray, preparations in which fully processed and C-terminally extended peptides occured are shown in orange, and preparations in which only the respective C-terminally extended peptide was detectable are shown in blue.

**Supporting Figure 2**: Examples of the LC-MS peptide analysis for two peptide families, Allatostatin A and FMRFa-like peptides. Identified peptides were aligned to the prepropeptide sequence. For both peptide families, fully processed and bioactive peptides are C-terminally amidated (red box); both spacer peptides and bioactive peptides were found. In *FM7h; hs-svr* control flies, only fully processed amidated bioactive peptides or mostly processed but not yet amidated (still carrying a C-terminal glycine amidation signal) are visible, with exception for AstA-1 (VERYAFGLa) which seems to have a weak C-terminal PC cleavage site. In *svr*^*PG33*^*;hs-svr* mutant flies, additional C-terminally extended peptides become detectable which still comprise the C-terminal PC cleavage sequence (KR, RR, R). Interestingly, a mutated dCPD leads sometimes to a skip of PC cleavage between some directly neighboured bioactive peptide sequences in both peptide families. Perhaps, intermediate dCPD action is required between two consecutive PC cleavage events in these cases.

**Supporting Figure 3**: Ratios R (Peak area_processed_/Peak area_unprocessed_), see Material and Methods) for *FM7h; hs-svr* control flies and *svr*^*PG33*^*;hs-svr* mutants. R>>1 suggests that more processed than unprocessed peptide is present, while R<<1 suggests that most peptide exists in its unprocessed form.

**Supporting Figure 4**: Phylogenetic analysis of M14 CPs by maximum likelihood method based on the JTT matrix-based model. The tree with the highest log likelihood (−253781,02) is shown. The percentage of trees in which the associated taxa clustered together is shown next to the branches. Initial tree(s) for the heuristic search were obtained automatically by applying Neighbor-Join and BioNJ algorithms to a matrix of pairwise distances estimated using a JTT model, and then selecting the topology with superior log likelihood value. The tree is drawn to scale, with branch lengths measured in the number of substitutions per site. The analysis involved 323 amino acid sequences, using insect CPA and CPB sequences as outgroup. All positions with less than 5% site coverage were eliminated. That is, fewer than 95% alignment gaps, missing data, and ambiguous bases were allowed at any position. There were a total of 1890 positions in the final dataset.

