## SupplementalMaterial for "*Drosophila* carboxypeptidase D (SILVER) is a key enzyme in neuropeptide processing required to maintain locomotor activity levels and survival rate"

**Supporting Information**

**Supporting table 1**: Primer pairs for RT-PCR

| Primer pair | sequence | melting temperature °C |
|---|---|---|
| primer 1 | forward: 5'-ATATCGGCATCAAGGGCCTG-3'<br>reverse: 5'-GCTGCTAATCGCCCTCAGAT -3' | 60.0<br>60.0 |
| primer 2 | forward: 5'-GTGTGCAATCCCATTACCGC-3'<br>reverse: 5'-CACCAAACAAAGCAGACGGG-3' | 59.9<br>60.0 |
| primer 3 | forward: 5'-CAAGCGCGCAATTAACCCTC-3'<br>reverse: 5'-AACTGGTGGACATGGCTCTG-3' | 60.5<br>59.8 |
| α-tubulin | forward: 5'-TCTGCGATTCGATGGTGCCCTTAAC-3'<br>reverse: 5'-GGATCGCACTTGACCATCTGGTTGGC-3' | 65.3<br>67.9 |

**Supporting Figure 1**: Summary of the qualitative MALDI-TOF MS peptide profiling in control (*FM7h; hs-svr*) and mutant (*svr*$^{PG33}$*;hs-svr*) flies. The total number of preparations with signals for the respective peptides is given inside the bars, the X-axis shows the relative % of detections. Preparations in which only the respective fully processed peptide was found are shown in gray, preparations in which fully processed and C-terminally extended peptides occured are shown in orange, and preparations in which only the respective C-terminally extended peptide was detectable are shown in blue.

**Supporting Figure 2**: Examples of the LC-MS peptide analysis for two peptide families, Allatostatin A and FMRFa-like peptides. Identified peptides were aligned to the prepropeptide sequence. For both peptide families, fully processed and bioactive peptides are C-terminally amidated (red box); both spacer peptides and bioactive peptides were found. In *FM7h; hs-svr* control flies, only fully processed amidated bioactive peptides or mostly processed but not yet amidated (still carrying a C-terminal glycine amidation signal) are visible, with exception for AstA-1 (VERYAFGLa) which seems to have a weak C-terminal PC cleavage site. In *svr*$^{PG33}$*;hs-svr* mutant flies, additional C-terminally extended peptides become detectable which still comprise the C-terminal PC cleavage sequence (KR, RR, R). Interestingly, a mutated dCPD leads sometimes to a skip of PC cleavage between some directly neighboured bioactive peptide sequences in both peptide families. Perhaps, intermediate dCPD action is required between two consecutive PC cleavage events in these cases.

**Supporting Figure 3**: Ratios R (Peak area$_{processed}$/Peak area$_{unprocessed}$), see Material and Methods) for *FM7h; hs-svr* control flies and *svr*$^{PG33}$*;hs-svr* mutants. R>>1 suggests that more processed than unprocessed peptide is present, while R<<1 suggests that most peptide exists in its unprocessed form.

**Supporting Figure 4**: Phylogenetic analysis of M14 CPs by maximum likelihood method based on the JTT matrix-based model. The tree with the highest log likelihood (-253781,02) is shown. The percentage of trees in which the associated taxa clustered together is shown next to the branches. Initial tree(s) for the heuristic search were obtained automatically by applying Neighbor-Join and BioNJ algorithms to a matrix of pairwise distances estimated using a JTT model, and then selecting the topology with superior log likelihood value. The tree is drawn to scale, with branch lengths measured in the number of substitutions per site. The analysis involved 323 amino acid sequences, using insect CPA and CPB sequences as outgroup. All positions with less than 5% site coverage were eliminated. That is, fewer than 95% alignment gaps, missing data, and ambiguous bases were allowed at any position. There were a total of 1890 positions in the final dataset.

SFig1

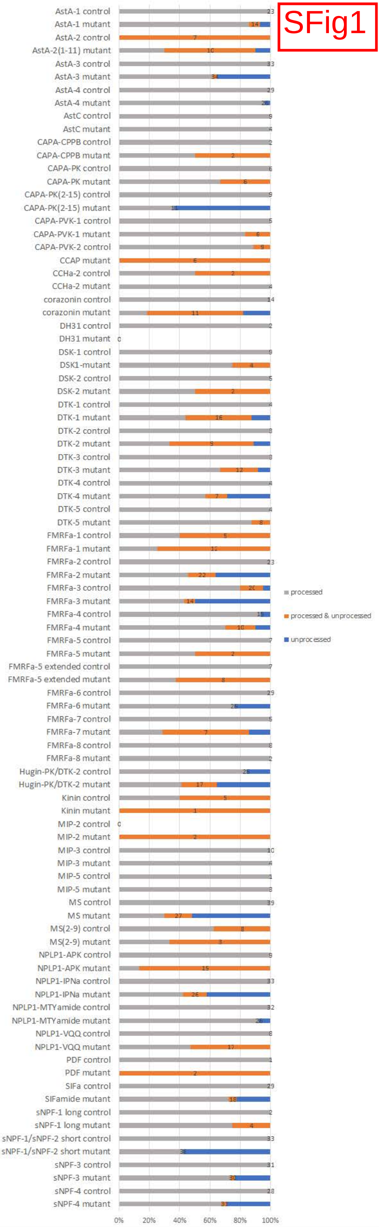
AstA-1 control
AstA-1 mutant
AstA-2 control
AstA-2(1-11) mutant
AstA-3 control
AstA-3 mutant
AstA-4 control
AstA-4 mutant
AstC control
AstC mutant
CAPA-CPPB control
CAPA-CPPB mutant
CAPA-PK control
CAPA-PK mutant
CAPA-PK(2-15) control
CAPA-PK(2-15) mutant
CAPA-PVK-1 control
CAPA-PVK-1 mutant
CAPA-PVK-2 control
CCAP mutant
CCHa-2 control
CCHa-2 mutant
corazonin control
corazonin mutant
DH31 control
DH31 mutant
DSK-1 control
DSK1-mutant
DSK-2 control
DSK-2 mutant
DTK-1 control
DTK-1 mutant
DTK-2 control
DTK-2 mutant
DTK-3 control
DTK-3 mutant
DTK-4 control
DTK-4 mutant
DTK-5 control
DTK-5 mutant
FMRFa-1 control
FMRFa-1 mutant
FMRFa-2 control
FMRFa-2 mutant
FMRFa-3 control
FMRFa-3 mutant
FMRFa-4 control
FMRFa-4 mutant
FMRFa-5 control
FMRFa-5 mutant
FMRFa-5 extended control
FMRFa-5 extended mutant
FMRFa-6 control
FMRFa-6 mutant
FMRFa-7 control
FMRFa-7 mutant
FMRFa-8 control
FMRFa-8 mutant
Hugin-PK/DTK-2 control
Hugin-PK/DTK-2 mutant
Kinin control
Kinin mutant
MIP-2 control
MIP-2 mutant
MIP-3 control
MIP-3 mutant
MIP-5 control
MIP-5 mutant
MS control
MS mutant
MS(2-9) control
MS(2-9) mutant
NPLP1-APK control
NPLP1-APK mutant
NPLP1-IPNa control
NPLP1-IPNa mutant
NPLP1-MTYamide control
NPLP1-MTYamide mutant
NPLP1-VQQ control
NPLP1-VQQ mutant
PDF control
PDF mutant
SIFa control
SIFamide mutant
sNPF-1 long control
sNPF-1 long mutant
sNPF-1/sNPF-2 short control
sNPF-1/sNPF-2 short mutant
sNPF-3 control
sNPF-3 mutant
sNPF-4 control
sNPF-4 mutant
0%
20%
40%
60%
80%
100%
processed
processed & unprocessed
unprocessed

SFig2

### Allatostatin A

#### *FM7h;hs-svr* run 1

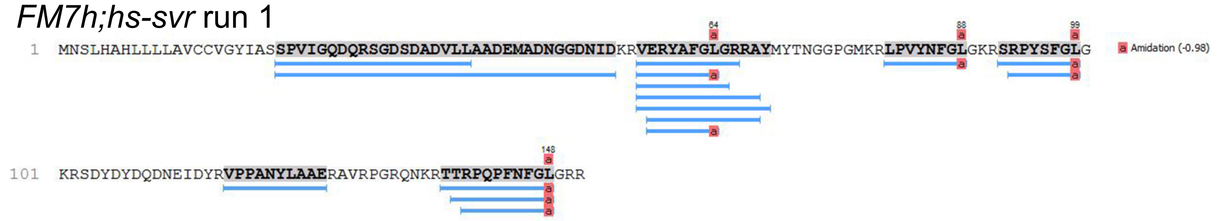


#### *svr*[PG33]*;hs-svr* run 2

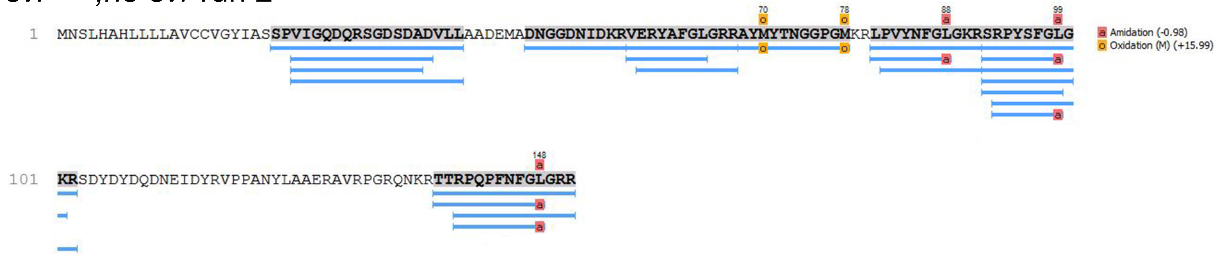


### FMRFa-like peptides

#### *FM7h;hs-svr* run 1

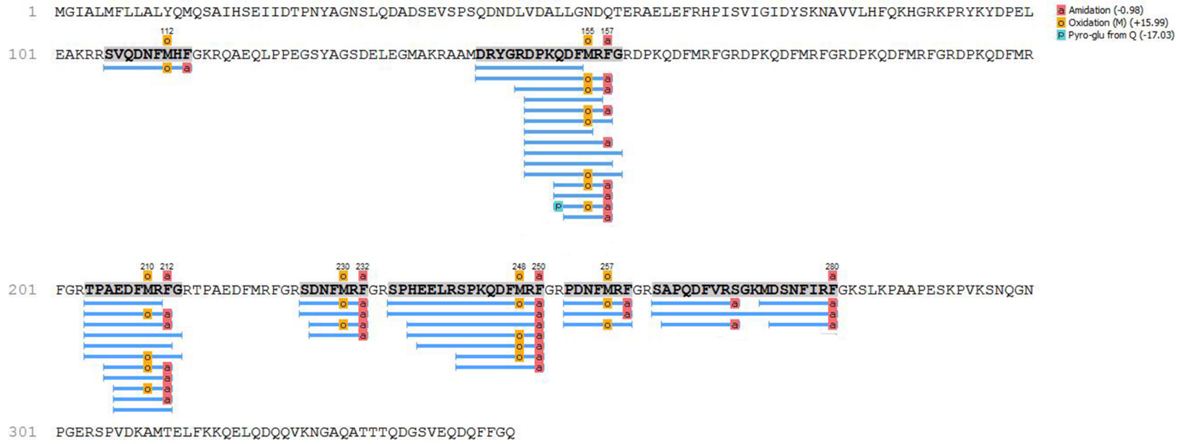


#### *svr*[PG33]*;hs-svr* run 3

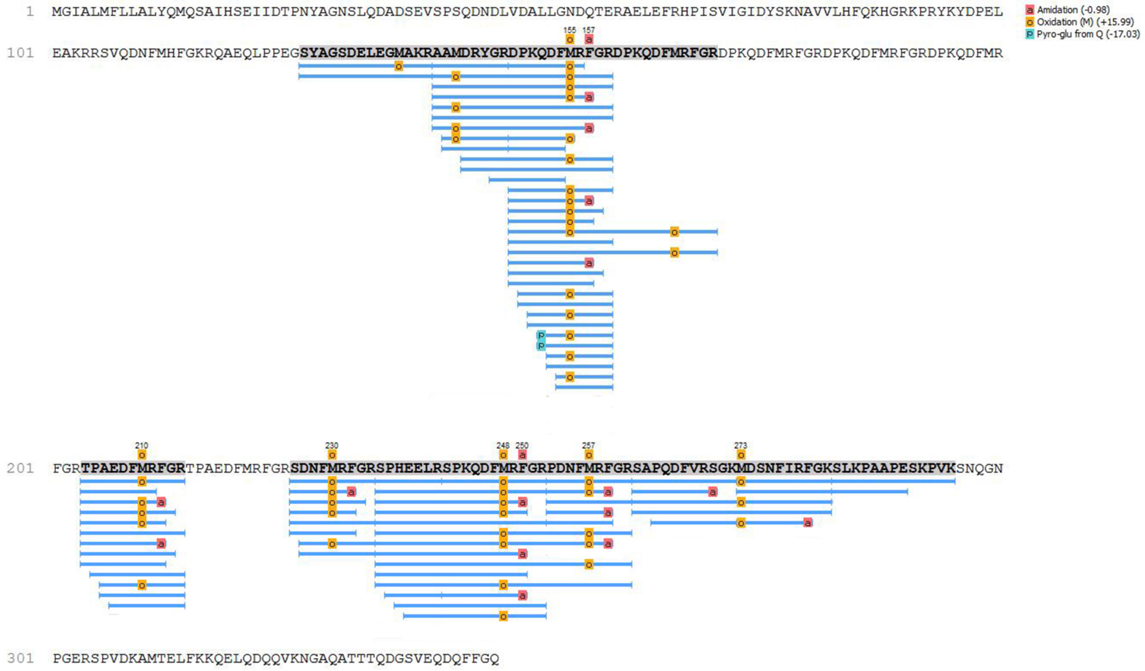

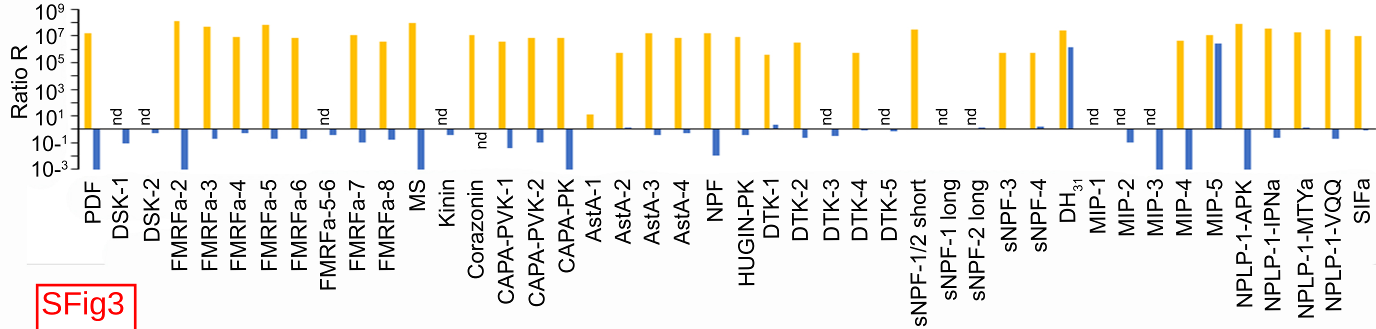

Ratio R
10^9
10^7
10^5
10^3
10^1
10-1
10-3
nd
PDF
DSK-1
DSK-2
FMRFa-2
FMRFa-3
FMRFa-4
FMRFa-5
FMRFa-6
FMRFa-5-6
FMRFa-7
FMRFa-8
MS
Kinin
Corazonin
CAPA-PVK-1
CAPA-PVK-2
CAPA-PK
AstA-1
AstA-2
AstA-3
AstA-4
NPF
HUGIN-PK
DTK-1
DTK-2
DTK-3
DTK-4
DTK-5
sNPF-1/2 short
sNPF-1 long
sNPF-2 long
sNPF-3
sNPF-4
DH31
MIP-1
MIP-2
MIP-3
MIP-4
MIP-5
NPLP-1-APK
NPLP-1-IPNa
NPLP-1-MTYa
NPLP-1-VQQ
SIFa

SFig3

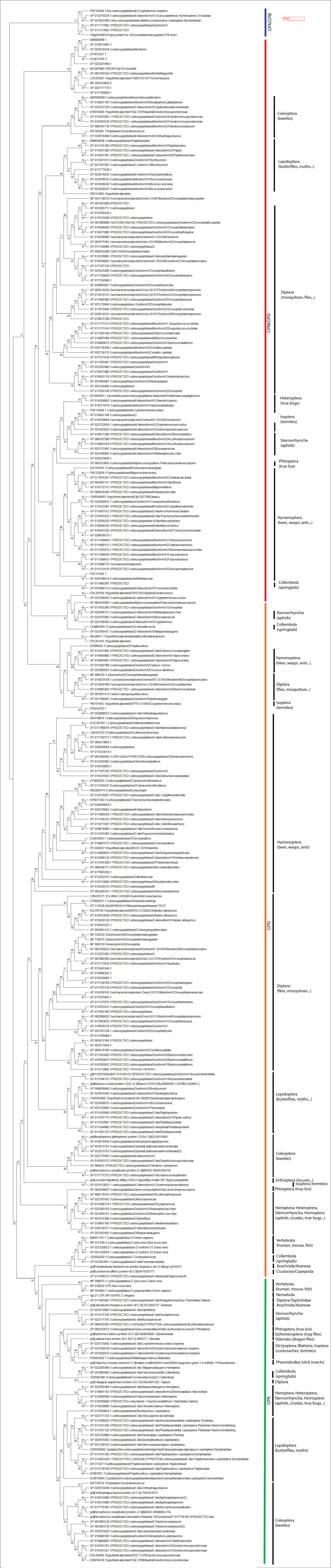

CPA/CPB
CPM/CPD
Coleoptera (beetles)
Lepidoptera (butterflies, moths...)
Diptera (mosquitoes, flies...)
Heteroptera (true bugs)
Isoptera (termites)
Sternorrhyncha (aphids)
Phtiraptera (true lice)
Hymenoptera (bees, wasps, ants...)
Collembola (springtails)
Sternorrhyncha (aphids)
Collembola (springtails)
Hymenoptera (bees, wasps, ants...)
Diptera (flies, mosquitoes...)
Isoptera (termites)
CPD
Hymenoptera (bees, wasps, ants)
Diptera (flies, mosquitoes...)
Lepidoptera (butterflies, moths...)
Coleoptera (beetles)
Orthoptera (locusts...)
Isoptera (termites)
Phtiraptera (true lice)
Hemiptera: Heteroptera, Sternorrhyncha, Homoptera (aphids, cicadas, true bugs...)
Vertebrata (human, mouse, fish)
Collembola (springtails)
Arachnida/Araneae
Crustacea/Copepoda
Vertebrata (human, mouse, fish)
Nematoda
Diptera/Tephritidae
Arachnida/Araneae
Sternorrhyncha (aphids)
Phtiraptera (true lice)
Ephemeroptera (may flies)
Odonata (dragon flies)
Dictyoptera: Blattaria, Isoptera (cockroaches, termites)
Phasmatodea (stick insects)
Collembola (springtails)
Diplura
Hemiptera: Heteroptera, Sternorrhyncha, Homoptera (aphids, cicadas, true bugs...)
Lepidoptera (butterflies, moths)
Coleoptera (beetles)
CPE
